# Transmission patterns of hyper-endemic multi-drug resistant *Klebsiella pneumoniae* in a Cambodian neonatal unit: a longitudinal study with whole genome sequencing

**DOI:** 10.1101/223701

**Authors:** Pieter W. Smit, Nicole Stoesser, Sreymom Pol, Esther van Kleef, Mathupanee Oonsivilai, Pisey Tan, Leakhena Neou, Claudia Turner, Paul Turner, Ben S Cooper

**Affiliations:** Centre for Tropical Medicine and Global Health, Nuffield Department of Medicine, University of Oxford, Old Road campus, Roosevelt Drive, Headington OX3 7FZ, UK; Mahidol Oxford Research Unit, Faculty of Tropical Medicine, Mahidol University, 3/F, 60th Anniversary Chalermprakiat Building, 420/6 Rajvithi Road, Bangkok 10400, Thailand; Nuffield Department of Clinical Medicine, University of Oxford, John Radcliffe Hospital (Level 5), Headley Way, Headington OX3 9DU, UK; John Radcliffe Hospital Microbiology Laboratory, John Radcliffe Hospital (Level 7), Headley Way, Headington OX3 9DU, UK; Cambodia-Oxford Medical Research Unit, Angkor Hospital for Children, Siem Reap, Combodia

**Keywords:** *Klebsiella pneumoniae*, multidrug-resistance, colonization, whole genome sequencing, neonatal unit

## Abstract

**Background:** *Klebsiella pneumoniae* is an important and increasing cause of life-threatening disease in hospitalised neonates. Third generation cephalosporin resistance (3GC-R) is frequently a marker of multi-drug resistance, and can complicate management of infections. 3GC-R *K. pneumoniae* is hyper-endemic in many developing country settings, but its epidemiology is poorly understood and prospective studies of endemic transmission are lacking. We aimed to determine the transmission dynamics of 3GC-R *K. pneumoniae* in a newly opened neonatal unit (NU) in Cambodia.

**Methods:** We performed a prospective longitudinal study between September and November 2013. Rectal swabs from 37 consented patients were collected upon NU admission and every three days thereafter. Morphologically different colonies from swabs growing cefpodoxime-resistant *K. pneumoniae* were selected for whole-genome sequencing (WGS).

**Results:** 32/37 (86%) patients screened positive for 3GC-R *K. pneumoniae* and 93 colonies from 119 swabs were sequenced. Isolates were resistant to a median of six (range 3-9) antimicrobials. WGS revealed high diversity; pairwise distances between isolates from the same patient were either 0-1 SNV or >1,000 SNVs; 19/32 colonized patients harboured *K. pneumoniae* colonies differing by >1000 SNVs. Diverse lineages accounted for 18 probable importations to the NU and nine probable transmission clusters involving 19/37 (51%) of screened patients. Median cluster size was 5 patients (range 3-9).

**Conclusions:** The epidemiology of 3GC-R *K. pneumoniae* was characterised by multiple introductions and a dense network of cross-infection, with half of screened neonates part of a transmission cluster. Efforts to reduce the 3GC-R *K. pneumoniae* disease burden should consider targeting both processes.

## Introduction

Multi-drug resistant (MDR) *Klebsiella pneumoniae* is an increasing problem worldwide and is associated with poor patient outcomes, particularly in vulnerable groups such as neonates^1–7^. Whole genome sequencing (WGS) of clinical isolates has demonstrated the importance of the clonal spread in of *K. pneumoniae* in outbreak settings^2,8^, and colonization is known to substantially increase the risk of infection^9,10^. The underlying carriage dynamics of MDR *K. pneumoniae* in normal circumstances in high risk populations, however, remain poorly defined and longitudinal carriage studies in neonatal intensive care units and high endemicity settings are lacking. Importantly, it is not clear from previous reports whether reported transmission links between patients identified by WGS in outbreak settings represent unusual occurrences related to sporadic lapses in infection control or, on the contrary, whether frequent but hidden patient-to-patient transmission is the norm.

In a prospective 12-month study in a newly opened neonatal unit (NU) in Cambodia, we described hyper-endemic colonization with highly drug-resistant *K. pneumoniae*^7^. Three-quarters of neonates in the unit were colonised with MDR *K. pneumoniae/oxytoca* and 2% (6/333) of these neonates developed hospital-acquired bacteraemia with a *K. pneumoniae* strain sharing the same antimicrobial susceptibility pattern as the colonising strain. Here we focus on the first two months of data from this study and use WGS to provide a high-resolution snapshot of the underlying transmission dynamics. Such genomic approaches have been used for investigating transmission dynamics in an adult ICU in a high income setting^10^, but have not been used to systematically investigate asymptomatic transmission in neonatal settings or in populations with a high level of endemic MDR *K. pneumoniae* infection. Longitudinal carriage studies are also lacking from developing country settings.

## Methods

The study took place in 2013 in a newly opened 11-bedded NU at the Angkor Hospital for Children, a non-governmental pediatric hospital in Siem Reap, Cambodia. The NU has two multi-patient rooms (one for intensive care and one for lower acuity patients; nurse:patient ratios over the study period were 0.63 and 0.40 respectively). There is a single isolation room. Angkor Hospital for Children has an active infection control programme, including bleach-based environmental cleaning and monitoring of staff hand-hygiene compliance^11^. Alcohol handrub is available by each bed and at ward entrances. Parents or guardians of neonates admitted to the NU during the study period were asked to provide consent. Rectal swabs were obtained from neonates entered into the study within 24 hours of admission and twice weekly until NU discharge. Seven environmental sites within the NU were sampled twice weekly (six sink plug holes which were sampled by swabbing around plug hole and the unit computer keyboard). Here we consider only *K. pneumoniae* isolates resistant to cefpodoxime (a marker for multidrug-resistance) derived from swabs taken over a seven week period following the opening of the NU. Patient swab results were not fed back to the NU clinicians.

Samples were processed and the extended spectrum beta-lactamase (ESBL)-producing phenotype defined as previously described^7^. In brief, swabs were inoculated onto MacConkey agar (Oxoid). Morphologically distinct cefpodoxime and/or imipenem resistant isolates, as observed by the laboratory technicians, were sub-cultured; species identification as *K. pneumoniae* was undertaken using a panel of biochemical tests. All isolates were tested for susceptibility to the following antimicrobials by the disk diffusion method: ampicillin, co-amoxiclav, ceftriaxone, ciprofloxacin, gentamicin, co-trimoxazole, ceftazidime, chloramphenicol, imipenem, cefpodoxime, and nitrofurantoin. Extended spectrum beta-lactamase (ESBL) production was determined using the double-disk method (cefotaxime +/-clavulanate and ceftazidime +/-clavulanate), following Clinical & Laboratory Standards Institute guidelines^12^. Strains were considered MDR if non-susceptible to ≥3 antimicrobial categories^13^. The first 96 morphologically different colonies from swabs growing cefpodoxime-resistant *K. pneumoniae* were selected for WGS. DNA was extracted from pure *K. pneumoniae* sub-cultures using the Fujifilm Quickgene system (Fujifilm, Tokyo, Japan) as per the manufacturer’s instructions, with an additional mechanical lysis step (FastPrep-24; MP Biomedicals, Santa Ana, CA, USA) following chemical lysis. DNA extracts were sequenced on the HiSeq 2500 platform, generating 150bp paired-end reads. Sequence data have been deposited in NCBI database (BioProject number: PRJNA395864).

To identify single nucleotide variants (SNVs), reads were mapped to the *K. pneumoniae* MGH78578 reference (GenBank: CP000647.1), and variants called as described previously^14^. De novo assemblies were generated using Velvet/Velvet optimiser^15^, with *in silico* multilocus sequence typing (MLST) performed using BLASTn to query the assemblies with reference alleles/sequence types registered in the Pasteur *K. pneumoniae* MLST database (http://bigsdb.pasteur.fr/klebsiella/klebsiella.html; 100% match over 100% length of reference sequence required to call an allele).

Phylogenetic trees were constructed using hierarchical clustering^16^ and ClonalFrameML^17^. Plasmid replicons were detected using PlasmidFinder^18^ (accessed 10 May 2016), using the SRST2 tool^19^. The ARG-ANNOT tool (accessed 23 May 2017) was used to detect known antibiotic-resistance genes^20^. Strains were defined as sequences with 0-1 SNV differences. Strain clusters were groups of different strains defined using an iterative procedure that placed two strains in the same cluster if they differed by ≤10 SNVs^21^. Patient clusters were defined as patients harbouring isolates of the same strain/strain cluster. Probable importation events were defined as a positive *K. pneumoniae* swab taken ≤48hrs after NU admission. Probable acquisition events were defined as a first positive swab taken >48 hours after NU admission together with initial negative swab. Plausible patient sources for acquisition events were considered to be neonates harbouring the same strain/strain cluster who were on the NU prior to the acquisition event and discharged no more than seven days prior to the acquiring patient’s admission date.

We used a permutation test to evaluate whether strains belonging to two sequence types (STs) had an increased likelihood of carrying the same plasmid or sharing phenotypic resistance to the same antibiotic if located within the same host (see Supplementary Material). We tested for associations between time since unit admission and the number of plasmid replicons, resistance genes and phenotypic resistances within a sample using random effects Poisson regression models (adjusting for within-host clustering effects). Statistical analysis was performed in R^22^.

The study was approved by the Angkor Hospital for Children Institutional Review Board and the Oxford Tropical Ethics Committee.

## Results

During the seven week period, 158 samples from 37 patients and seven environmental locations were collected. Of these, 79 samples (from 32 patients and three environmental sites [all sinks]) were cefpodoxime-resistant *K. pneumoniae* culture-positive, generating 105 MDR *K. pneumoniae* isolates (Figure 1). The first 96 of these isolates were selected for sequencing.

**Figure 1.**
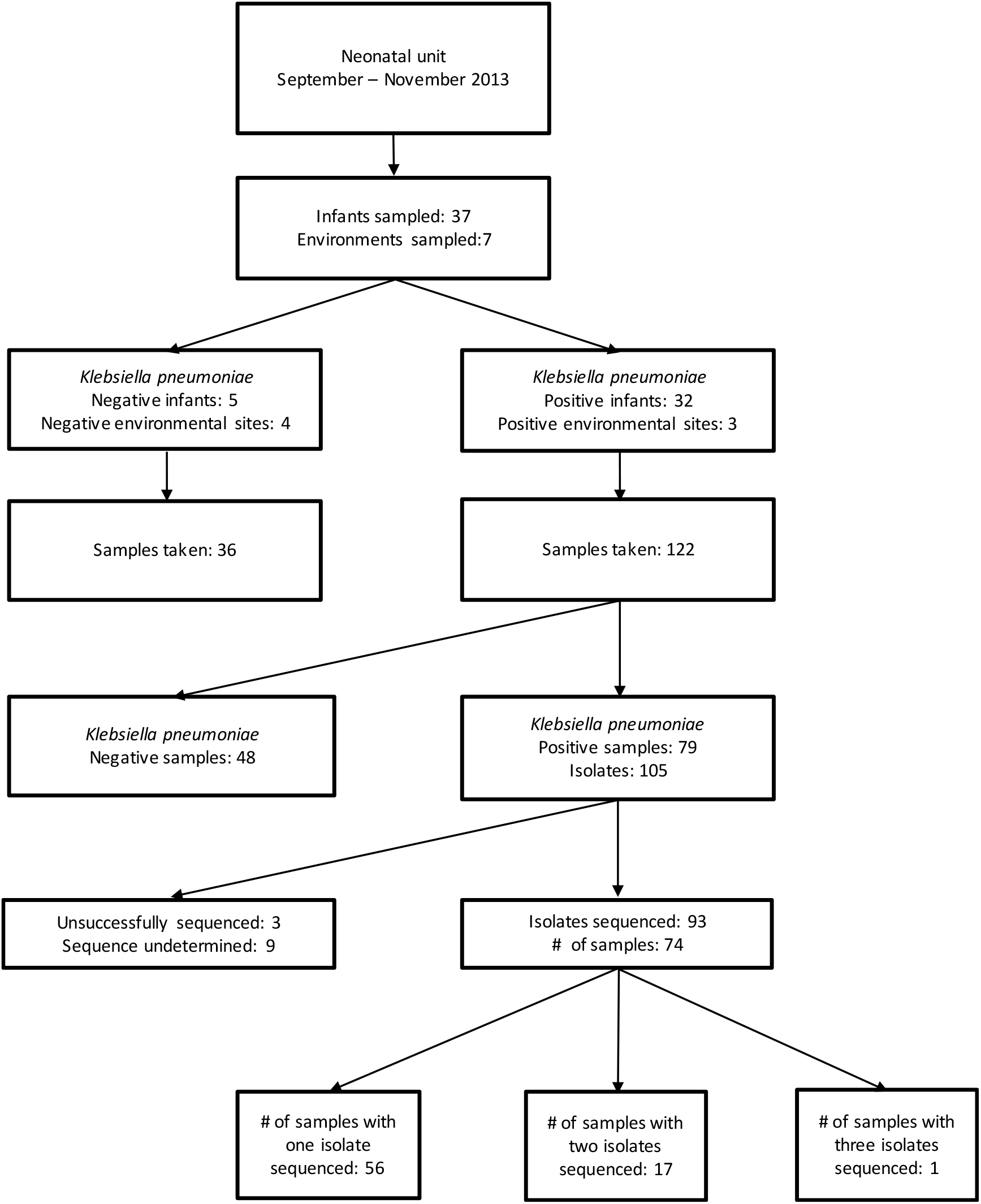
Flow diagram of study sampling strategy.

All 32 colonised patients were born outside the NU, with 16 born at a hospital, 13 at a health centre and three at home. None of these patients developed culture-proven *K. pneumoniae* invasive infection during the study timeframe.

Seventeen neonates (46% of those screened, 53% of those who were ever carrying MDR *K. pneumoniae)* were found to be colonised within 24 hours of NU admission, eight had their first positive sample at 1–3 days after admission and seven had a first positive sample 3–14 days after NU admission (median: four days). Of the seven environmental sites that were screened, three sink basins located in the isolation room, dirty utility, and milk kitchen room were intermittently positive. The first sink became positive 12 days after the first patient arrived.

### Antimicrobial resistance

Of 93 successfully sequenced isolates, all were MDR, with phenotypic resistance to a median of six of the 11 antibiotics tested (Figure 2); 17 isolates were resistant to ≥8 antibiotics and 22 were resistant to all antibiotic classes tested except nitrofurantoin and quinolones. Only one isolate did not have an ESBL phenotype. No isolates were resistant to imipenem. Amongst the 18 samples from which multiple colonies were selected and successfully sequenced, all morphologically distinguishable colonies had a different antimicrobial resistance profile.

**Figure 2.**
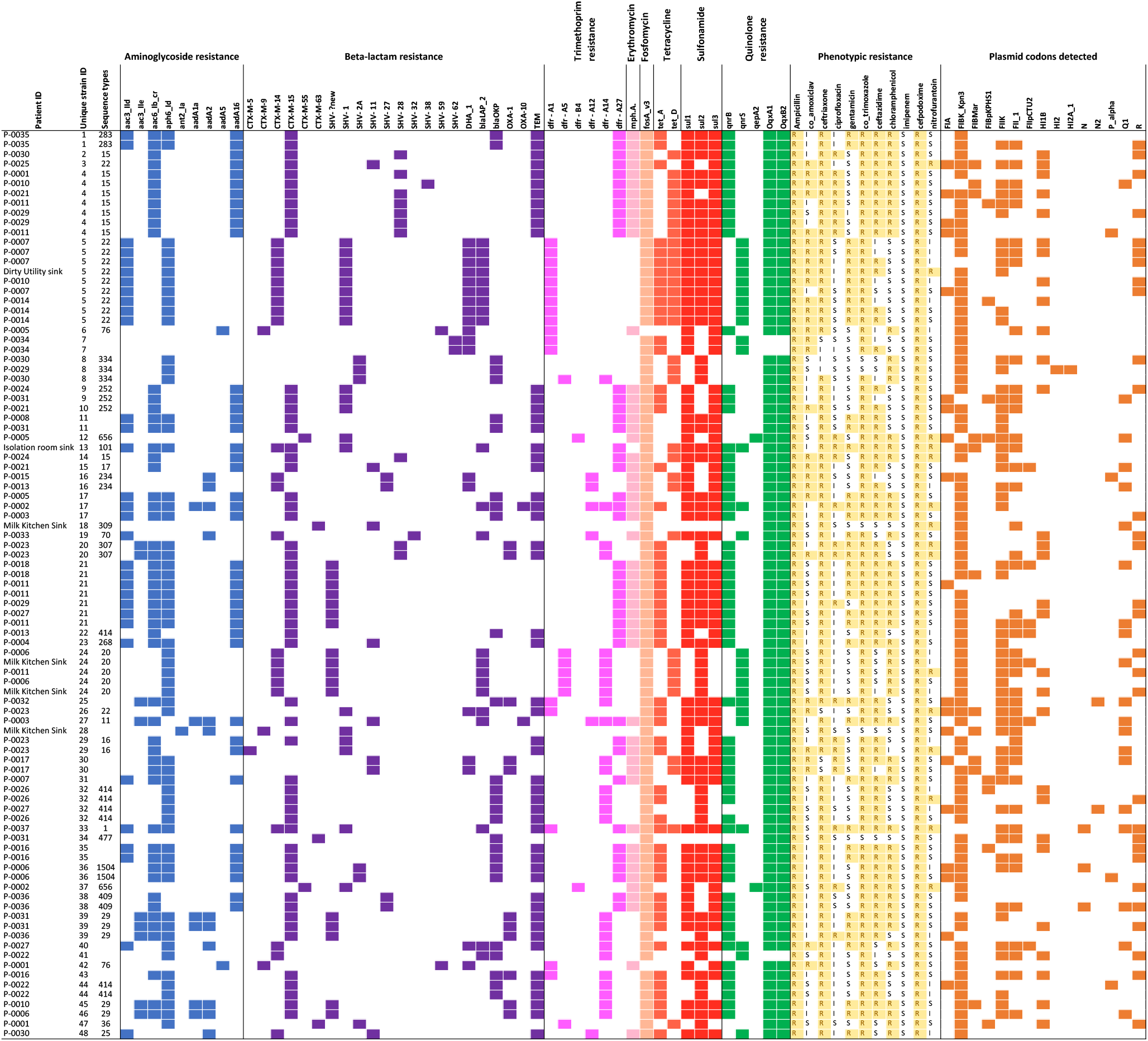
Overview of all isolates with colored blocks representing the presence of antimicrobial resistance genes, phenotypic resistance, or plasmid replicons. The list is sorted per unique strain ID (cut-off used: 0-1 single nucleotide variant difference), highlighting variations in phenotypic resistance and plasmid replicons detected among genetically identical strains

A total of 21 resistance genes (49 allelic variants) were detected, and isolates harboured an average of 14 allelic variants of resistance genes (range: 5–24). The relationship between phenotypic resistance and the presence of known resistance genes is shown in Supplementary Figures 1 and 2. A histogram showing the frequency of different phenotypic antimicrobial resistance combinations is shown in Supplementary Figure 3.

**Figure 3.**
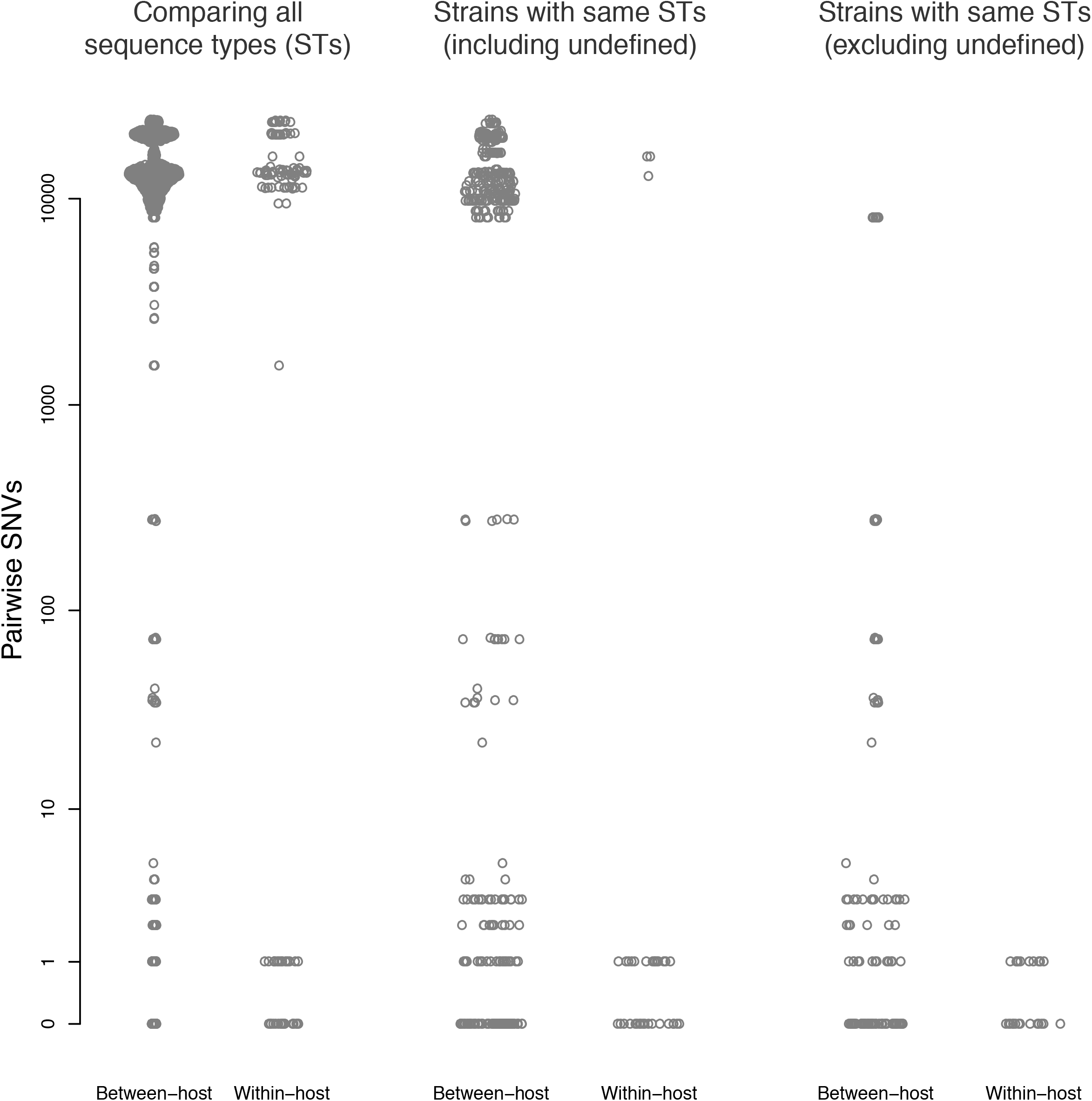
Pairwise single nucleotide variants (SNVs) between sequences. Results are shown for all pairwise comparisons (left), excluding pairs with different STs (centre), and excluding pairs with different STs or undefined ST (right). Points are jittered in proportion to the kernel density estimate.

### Whole genome sequencing

Sixty-one strains were identified amongst the 93 successfully sequenced colonies. Thirteen of these were represented by ≥1 isolate, and 9 strains were found in two or more patients. WGS revealed high genetic diversity among the sequenced isolates (median 13,580 SNVs, range 0-24,052). Twenty-five different STs were identified from 69 isolates where ST could be established; 10 of these STs were identified only once.

### Within- and between-host diversity

Thirty patients (94%) and three environmental sites had more than one isolate sequenced. Amongst these, within-host pairwise distances were either 0–1 or >1,000 SNVs (Figure 3). There were 45 sequences that differed ≤1 SNV from other sequenced isolates from the same patient, and 55 sequences that differed by ≤10 SNVs from sequences from a different patient.

Nine strain clusters were detected, and identified amongst 19/32 colonised patients in patient clusters ranging in size from three to nine patients (median 5). Ten patients (31%) were part of multiple patient clusters (Figure 4). In a sensitivity analysis where we varied the SNV threshold in the strain cluster definition from 10 to either 5 or 20 we identified identical strain and patient clusters.

**Figure 4.**
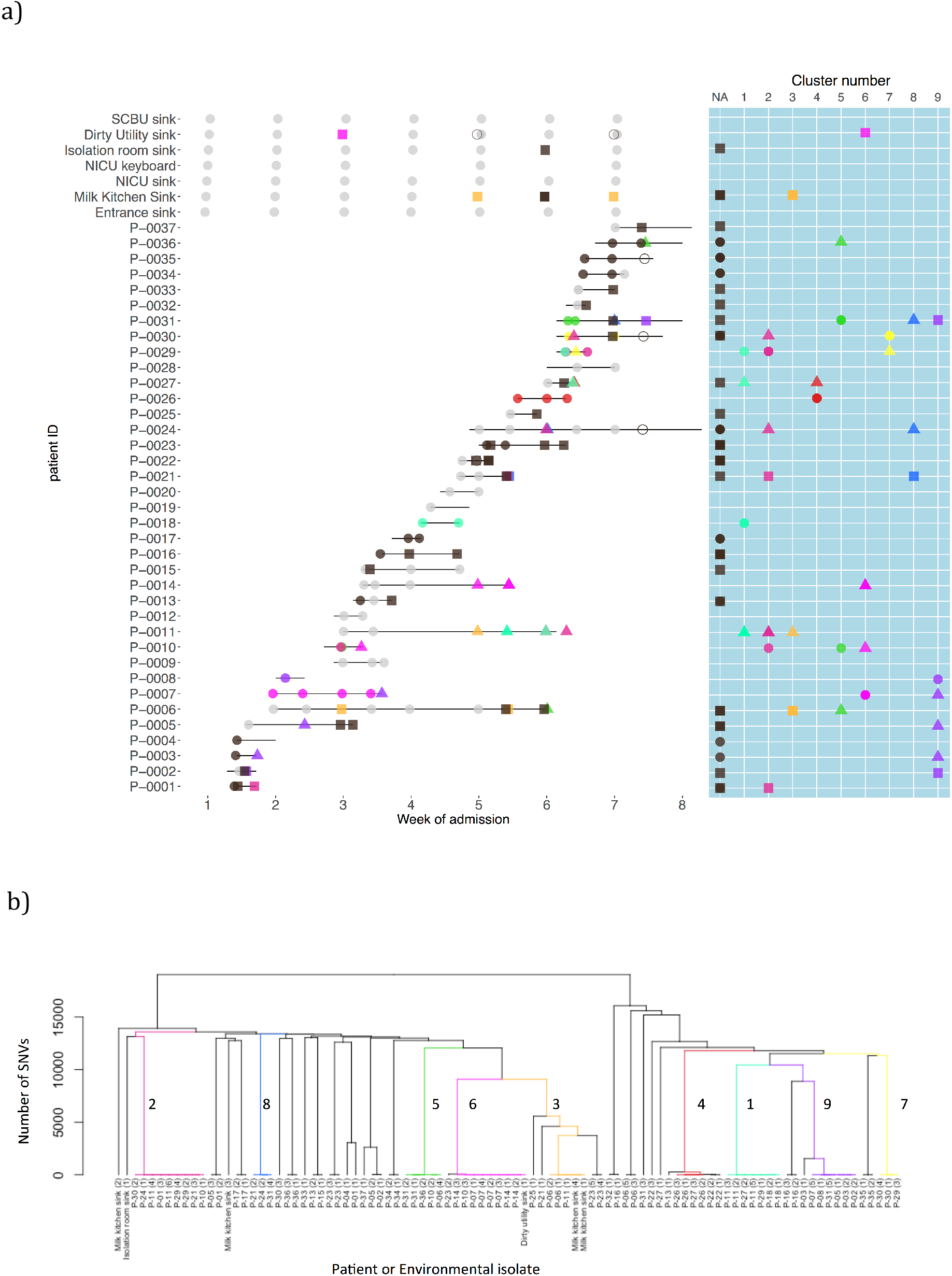
Figure 4. Patient carriage and environmental contamination with clustered and unclustered multidrug-resistant (MDR) *K. pneumoniae* strains and associated phylogentic tree, a) shows the timeline for individual neonatal admissions (lines) and the identification of clustered isolates. Light gray dots represent negative samples, dark gray indicates samples positive for MDR *K. pneumoniae* but with non-clustered strains. Other colors depict genetically-defined strain clusters. Shapes correspond to inferred importation/acquisition events; overlapping shapes indicate that different strains were detected from the same patient or environmental sample. Where the same colored shape is represented consecutively, this represents persistent carriage with a strain that was either imported (consecutive circles) or acquired on the neonatal unit (consecutive squares or triangles), b) Hierarchical phylogenetic tree of the isolates included in this study. Colors depict isolates from the same genetically-defined strain clusters as shown in a).

### Strain importations and acquisitions

Among the 32 *K. pneumoniae* positive patients, 18 (56%) represented probable importations. However, 11 of these patients (61%) also showed evidence of acquiring additional *K. pneumoniae* strains during their NU admission. There were 52 such possible acquisitions (defined by a first positive swab for a strain at least 48 hours after unit admission preceded by at least one negative swab for that strain/strain cluster) which occurred in 25 patients. In 22 of these 52 possible acquisitions a plausible source was identified. Environmental sources were identified as plausible sources for acquisitions in 2/9 clusters, though in both cases other patients were also plausible sources.

### Plasmids

We identified 15 known plasmid replicon sequences. All isolates were positive for at least one plasmid replicon, with an average of 3.4 replicon types per isolate (range 1–6) (Figure 2, Supplementary Table 1). Identical strains which carried the same resistance genes, had different plasmids identified. Neither of the plasmid-mediated colistin-resistance genes, *mcr-1* and *mcr-2*, were detected.

We found no evidence of systematic changes in the number of distinct plasmids, the number of resistance genes, or the number of antibiotics to which isolates were phenotypically resistant over the course of a patient’s stay (Supplementary Figure 4, Supplementary Table 2). We found no evidence of an association between the number of distinct plasmid replicons detected and either the number of resistance genes identified or the number of antibiotics to which isolates were phenotypically resistant, though (as expected) there was a positive correlation between the number of resistance genes and the phenotypic resistance count (Supplementary Figure 5). We found no evidence that strains from two STs were more likely to harbour the same plasmid if located within the same patient (Supplementary Table 3). Similarly, with the exception of nitrofurantoin, there was no statistical evidence of within-host transfer of phenotypic resistance between different STs (Supplementary Table 4).

## Discussion

The high frequency with which MDR *K. pneumoniae* isolates differing by ≤10 SNVs were recovered from different neonates together with the timing of probable acquisition events suggested frequent cross-infection between neonates and a high background rate of importation of new strains (almost half of the neonates were colonised on admission; 32/37 (86%) were carriers at some point). These apparently high transmission rates occurred despite relatively good levels of infection control and antibiotic stewardship^11^. Individuals were commonly involved in multiple strain clusters and the high prevalence of co-infection with genetically distant lineages of MDR *K. pneumoniae* points to a complex epidemiology that differs substantially from both that found in prospective WGS carriage studies of *K. pneumoniae* in a high-income adult intensive care unit setting^10^, and that suggested by older carriage studies in high income neonatal intensive care units^23,24^. Because the decision to perform this study was not motivated by an unusual clusters of cases, the results strongly suggest that such high rates of transmission of asymptomatic infection may be typical in neonatal units where MDR *K. pneumoniae* is endemic.

Quantifying the role of environmental contamination in transmission within hospital units is difficult, as presence of contamination does not necessarily imply such contamination is important for transmission. Randomised trials (that reduce such contamination) are needed to definitively establish a causal link and quantify the effects on transmission. However, contaminated sinks have been associated with previous outbreaks and it has been suggested that more intensive cleaning protocols might be needed^25^. In our study, environmental sites showed only intermittent contamination, although three out of six sinks screened positive on at least one occasion. However, none of these sites showed persistent colonisation with a single lineage and rates for positivity were low suggesting that cleaning protocols were effective and that the environment may have played a limited role in the transmission dynamics.

Strengths of this work include the use of systematic and high frequency screening of most neonates in the unit, highly discriminatory typing, and the sequencing of multiple morphologically distinct colonies grown from the same swabs. Without these three approaches we would have had a far more limited view of the complex transmission network.

A limitation was the inability to associate specific resistance genes with specific plasmids. This cannot usually be done reliably without fully resolving plasmid structures using long-read sequencing^26^. Also, by limiting our analysis to cefpodoxime-resistant *K. pneumoniae* we have only a partial epidemiological picture. It is possible, for example, that some of the apparent acquisition events of MDR *K. pneumoniae* where no plausible source was identified may have resulted from the transfer of mobile genetic elements between patients conferring cefpodoxime-resistance to previously cefpodoxime-sensitive strains, as a result of transfer from other colonising species which were not surveyed, or as a result of failure to completely characterise within-host diversity. In relation to the latter, morphology may not be the most sensitive way to distinguish within-host diversity; multiple picks, irrespective of colonial morphology, may uncover significant additional diversity^27^.

There are three key findings from this study that have important implications for attempts to control the spread of MDR *K. pneumoniae*. The first is to suggest that super-infection with diverse lineages of *K. pneumoniae* might be the norm rather than exception in neonatal populations, highlighting the importance of considering the carrier state and taking more than one sample per patient, and investigating more than one colony within a sample culture, when trying to understand transmission chains of *K. pneumoniae* (similar considerations have been shown to be important for ESBL-producing *Escherichia coli^21^)*. The second is to highlight the high degree of cross-infection in the NU despite good infection control and antibiotic stewardship practices. The third is to show that high numbers of infants are likely to have be already colonized when admitted to the unit, suggesting acquisition in the community may be important in this population. Together these observations highlight the scale of the challenge in halting the spread of MDR *K. pneumoniae* in endemic settings; our findings emphasise the importance and difficulty of both finding effective interventions to impede its nosocomial spread as well as measures to reduce the chance that infants are carrying this organism when admitted to hospital. Currently we are lacking strategies of proven effectiveness to do either.

### Funding

This study was supported by the UK Medical Research Council and Department for International Development [grant number MR/K006924/1 to B.S.C.], by the Wellcome Trust as part of the Wellcome Trust-Mahidol University-Oxford Tropical Medicine Research Programme [grant number 106698/Z/14/Z], and by the NIHR Oxford Biomedical Research Centre. NS is supported by a Public Health England/University of Oxford Clinical Lectureship.

## Acknowledgments

We would like to thank all staff in the neonatal unit at the Angkor Hospital for Children.

## References

1. Stapleton PJ, Murphy M, McCallion N, Brennan M, Cunney R, Drew RJ. Outbreaks of extended spectrum beta-lactamase-producing Enterobacteriaceae in neonatal intensive care units: a systematic review. Arch Dis Child Fetal Neonatal Ed, 2016; 101: F72–8.

2. Snitkin ES, Zelazny AM, Thomas PJ, et al. Tracking a hospital outbreak of carbapenem-resistant Klebsiella pneumoniae with whole-genome sequencing. Sci Transl Med, 2012; 4: 148ra116.

3. Munoz-Price LS, Poirel L, Bonomo RA, et al. Clinical epidemiology of the global expansion of Klebsiella pneumoniae carbapenemases. Lancet Infect Dis, 2013; 13: 785–796.

4. Falade AG, Ayede AI. Epidemiology, aetiology and management of childhood acute community-acquired pneumonia in developing countries – a review. Afr J Med Med Sci, 2011; 40: 293–308.

5. Hendrik TC, Voor In’t Holt AF, Vos MC. Clinical and molecular epidemiology of extended-spectrum beta-lactamase-producing Klebsiella spp.: a systematic review and metaanalyses. PLoS One, 2015; 10: e0140754.

6. Naas T, Cuzon G, Robinson AL, et al. Neonatal infections with multidrug-resistant ESBL-producing E. cloacae and K. pneumoniae in Neonatal Units of two different Hospitals in Antananarivo, Madagascar. BMC Infect Dis, 2016; 16: 275.

7. Turner P, Pol S, Soeng S, et al. High Prevalence of antimicrobial resistant Gram negative colonization in hospitalized Cambodian infants. Pediatr Infect Dis J, 2016;35: 856–861.

8. Stoesser N, Giess A, Batty EM, et al. Genome sequencing of an extended series of NDM-producing Klebsiella pneumoniae isolates from neonatal infections in a Nepali hospital characterizes the extent of community-versus hospital-associated transmission in an endemic setting. Antimicrob Agents Chemother, 2014; 58: 7347–7357.

9. Martin RM, Cao J, Brisse S, et al. Molecular epidemiology of colonizing and infecting isolates of Klebsiella pneumoniae. mSphere, 2016; 1: e00261–16.

10. Gorrie CL, Mirceta M, Wick RR, et al. Gastrointestinal carriage is a major reservoir of K. pneumoniae infection in intensive care patients. Clin Infect Dis 2017; 65: 208–215.

11. Stoesser N, Emary K, Soklin S, et al. The value of intermittent point-prevalence surveys ofhealthcare-associated infections fo evaluating infection control interventions at angkor hospital for children, siem reap, cambodia. Trans R Soc Trop Med Hyg, 2013; 107: 248–253.

12. CLSI. Performance Standards for Antimicrobial Susceptibility Testing; Seventeenth Informational Supplement. CLSI Document M100-S23. Wayne, PA: Clinical and Laboratory Standards Institute; 2013.

13. Magiorakos AP, Srinivasan A, Carey RB, et al. Multidrug-resistant, extensively drug-resistant and pandrug-resistant bacteria: An international expert proposal for interim standard definitions for acquired resistance. Clin Microbiol Infect 2012; 18: 268–281.

14. Stoesser N, Xayaheuang S, Vongsouvath M, et al. Colonization with Enterobacteriaceae producing ESBLs in children attending pre-school childcare facilities in the Lao People’s Democratic Republic. J Antimicrob Chemother, 2015; 70:1893–1897.

15. Zerbino DR, BirneyE. Velvet: Algorithms for de novo short read assembly using de Bruijn graphs. Genome Res, 2008; 18: 821–829.

16. Müllner D. fastcluster: Fast hierarchical, agglomerative clustering routines for R and Python. J Stat Softw, 2013; 53:1–18.

17. Didelot X, Wilson DJ. ClonalFrameML: Efficient inference of recombination in whole bacterial genomes. PLoS Comput Biol, 2015; 11:1–18.

18. Carattoli A, Zankari E, García-Fernández A, et al. In silico detection and typing of plasmids using PlasmidFinder and plasmid multilocus sequence typing. Antimicrob Agents Chemother, 2014; 58: 3895–3903.

19. Inouye M, Dashnow H, Raven L-A, et al. SRST2: Rapid genomic surveillance for public health and hospital microbiology labs. Genome Med, 2014; 6: 90.

20. Gupta SK, Padmanabhan BR, Diene SM, et al. ARG-annot, a new bioinformatic tool to discover antibiotic resistance genes in bacterial genomes. Antimicrob Agents Chemother, 2014; 58: 212–220.

21. Tong S, Holden MT, Nickerson EK, et al. Genome sequencing defines phylogeny and spread of methicillin-resistant Staphylococcus aureus in a high transmission setting. Genome Res, 2015; 25:111–118.

22. R Core Team. R: A Language and Environment for Statistical Computing. Available at https://www.R-project.org/. Accessed 1 October 2017.

23. Toltzis P, Dul MJ, Hoyen C, et al. Molecular epidemiology of antibiotic-resistant gram-negative bacilli in a neonatal intensive care unit during a nonoutbreak period. Pediatrics, 2001; 108: 1143–1148.

24. Goldmann DA, Leclair J, Macone A. Bacterial colonization of neonates admitted to an intensive care environment. J Pediatr, 1978; 93: 288–193.

25. Lowe C, Willey B, O’Shaughnessy A, et al. Outbreak of extended-spectrum beta-lactamase-producing Klebsiella oxytoca infections associated with contaminated handwashing sinks. Emerg Infect Dis 2012; 18:1242–1247.

26. Holt KE, Wertheim H, Zadoks RN, et al. Genomic analysis of diversity, population structure, virulence, and antimicrobial resistance in Klebsiella pneumoniae, an urgent threat to public health. Proc Natl Acad Sci, 2015; 112: E3574–E3581.

27. Stoesser N, Sheppard AE, Moore CE, et al. Extensive within-host diversity in fecally carried extended-spectrum-beta-lactamase-producing Escherichia coli Isolates: Implications for Transmission Analyses. J Clin Microbiol, 2015; 53: 2122–2131.

